# When Multimodal Fusion Fails: Contrastive Alignment as a Necessary Stabilizer for TCR–Peptide Binding Prediction

**DOI:** 10.64898/2026.03.31.715453

**Authors:** Cong Qi, Wenbo Wang, Hanzhang Fang, Zhi Wei

## Abstract

Multimodal learning is commonly assumed to improve predictive performance, yet in biological applications auxiliary modalities are often imperfect and can degrade learning if fused naively. We investigate this problem in TCR–peptide binding prediction, where sequence embeddings from pretrained protein language models are strong and transferable, but structure-derived residue graphs are built from predicted folds and heuristic discretization. In this setting, structural views can be noisy, inconsistent, and difficult to optimize jointly with sequence features. We introduce TRACE, a lightweight multimodal framework that encodes each entity (TCR and peptide) with parallel sequence and graph towers, then applies CLIP-style intra-entity contrastive alignment before interaction modeling. The alignment objective regularizes representation geometry by encouraging modality consistency for the same biological entity, thereby preventing unstable graph signals from dominating fusion. Across protocol-aware TCHard RN evaluations, naive sequence+graph fusion frequently underperforms a sequence-only baseline and can collapse toward near-random behavior. In contrast, TRACE consistently restores and improves performance. Controlled noise and supervision sweeps show that these gains persist under increasing graph corruption and positive-label scarcity, indicating that alignment is especially important when training conditions are hard. Our results challenge the assumption that adding modalities is inherently beneficial. Instead, they highlight a central principle for robust multimodal bioinformatics: performance depends not only on what modalities are used, but on how their interaction is constrained during optimization. TRACE provides a simple and general recipe for leveraging imperfect structural information without sacrificing stability.

## 1 Introduction

Multimodal fusion is often assumed to help, yet when one modality is unstable or low-quality, naive fusion can hurt performance. Without constraints, noisy inputs can dominate gradients and distort representations learned from stronger modalities. This failure mode is easy to overlook because it may only appear under distribution shift or label scarcity.

Protein tasks provide a concrete example. Structure offers a powerful inductive bias, but practical pipelines often depend on predicted folds and heuristic graph construction, which are intrinsically noisy [Jumper et al., 2021]. As a result, simply combining sequence embeddings with structure-derived graphs can yield limited or inconsistent gains in protein representation learning [Kalifa et al., 2025].

At the same time, pretrained protein language models provide strong sequence representations that transfer well in low-data settings. Yet binding is driven by residue-level interactions, and pooled sequence embeddings alone can miss local geometric cues that structure could provide. This creates a tension: structure is informative but noisy, and sequence is reliable but coarse. A useful multimodal method must therefore leverage structural bias without letting structural noise override the sequence signal.

We study this issue in TCR–peptide binding prediction. T cell receptors recognize peptide antigens presented by major histocompatibility complex (pMHC) molecules, enabling applications such as neoantigen selection, vaccine design, and TCR engineering. In the supervised setting, the goal is to predict binding given a TCR (here the *β*-chain CDR3) and a peptide sequence. The task is challenging due to sparse labels, high TCR diversity, and protocol sensitivity: models that appear strong on random splits often fail to generalize to unseen epitopes or altered negative distributions [Deng et al., 2023, Dens et al., 2023, Castorina et al., 2025, Wang et al., 2025].

Recent predictors benefit from strong sequence encoders and interaction heads, but they often rely on the supervised objective to arbitrate between modalities. Under hard negatives or distribution shift, this can allow noisy structural signals to dominate, masking true residue-level specificity. These conditions motivate explicit constraints that regulate cross-modal interaction rather than assuming that adding a modality will help.

Early deep-learning approaches to TCR–peptide or TCR–pMHC binding prediction model TCR and peptide sequences directly, including convolutional architectures such as NetTCR [Jurtz et al., 2018] and subsequent extensions that incorporate paired chains and refined training strategies [Montemurro et al., 2021]. Beyond convolutional models, several works adopt attention-based or recurrent architectures and metric-learning formulations to capture epitope specificity and repertoire-level patterns [Weber et al., 2021, Bi et al., 2022, Sidhom et al., 2018, Jokinen et al., 2019]. More recent predictors continue to explore modeling choices for generalization and transfer, including epiTCR-style frameworks and related analyses [Pham et al., 2023, Kim et al., 2023, Jiang et al., 2022]. A recurring theme across these methods is the tension between expressiveness and generalization. While higher-capacity interaction modules can improve in-distribution accuracy, reported performance is often highly sensitive to the choice of split protocol and negative sampling strategy. Models that appear strong under random splits frequently fail to generalize to unseen epitopes or altered negative distributions [Deng et al., 2023, Dens et al., 2023, Castorina et al., 2025]. These observations motivate evaluation protocols that explicitly stress distribution shift and few-shot generalization, such as TCHard-style splits, and the use of threshold-free metrics and calibration analyses.

Several works incorporate additional biological context, such as MHC alleles, to improve binding specificity prediction [Springer et al., 2021, Long et al., 2025]. Others introduce structure-derived features or residue-level representations to capture local interaction patterns that are not accessible from pooled sequence embeddings alone [Li et al., 2024, Montemurro et al., 2021]. In this line of work, structure is typically obtained from predicted folds or contact maps and encoded as graphs, providing an explicit neighborhood operator for residue-level aggregation [Jumper et al., 2021]. More broadly, multimodal modeling has become an active direction in protein representation learning. While integrating sequence and structure information is intuitively appealing, prior attempts at static or one-shot fusion have often shown limited benefits over strong single-modality models, partly due to the loss of structural information and sensitivity to noisy predicted structures [Kalifa et al., 2025]. These findings suggest that multimodal performance is not determined solely by the presence of additional modalities, but by how their interaction is regulated during training. Alignment-based multimodal frameworks have therefore emerged as a principled alternative, emphasizing coordination between modalities rather than static feature concatenation [Bolouri et al., 2025, Flöge et al., 2025]. Although developed at the protein representation level, these methods provide an important methodological insight for TCR–peptide binding prediction.

Contrastive objectives such as InfoNCE-style losses are widely used to align multiple views of the same instance and to stabilize representation learning across modalities [Radford et al., 2021]. In parallel, graph neural networks offer a flexible mechanism for neighborhood aggregation and message passing in structured domains [Kipf and Welling, 2017, Veličković et al., 2018, Gilmer et al., 2017, Hamilton et al., 2017]. COATI [Kaufman et al., 2024] employs contrastive learning to align textual (SMILES) and 3D molecular representations, yielding a shared embedding space that supports downstream regression and molecular generation.

To address noisy multimodal fusion, we propose *TRACE* (*TCR Robust Alignment via Contrastive Encoding*), a lightweight framework that encodes each entity with parallel sequence and residue-graph towers and aligns them using an *intra-entity* contrastive objective [Oord et al., 2018, Chen et al., 2020, Radford et al., 2021]. Alignment regularizes how modalities interact and prevents the structural view from destabilizing learning when it is unreliable.

On protocol-aware TCHard-style splits, we show that naive sequence-plus-graph fusion underperforms a sequence-only baseline, while intra-entity alignment consistently restores and improves performance. These results emphasize a simple principle: in TCR–peptide binding prediction, how modalities are integrated matters more than how many are used.

## 2 Methods

### 2.1 Problem setup

We consider supervised binary classification for TCR–peptide binding. Each sample is a triple (*t, p, y*), where *t* is a TCR*β* CDR3 amino-acid sequence, *p* is a peptide sequence, and *y* ∈ {0, 1} indicates binding. For each entity (TCR or peptide), we use two modalities:

- **Sequence representations**. A pretrained protein language model provides a global sequence embedding **s** ∈ ℝ^*ds*^ .
- **Residue graphs**. A residue-level graph *G* = (*V, E*) is derived from a predicted fold. Nodes correspond to residues with node features **X** ∈ℝ^|*V* |×*dx*^ and edges encode sequence adjacency and spatial proximity. Because structures are predicted (not experimentally resolved) and discretized via heuristics, this modality can exhibit uncertainty or inconsistency across samples.

### 2.2 Graph construction from predicted folds

We predict 3D structure with ESMFold and retain C*α* atoms to represent residues. Each residue becomes a node with a 20D one-hot amino-acid identity feature. We add (i) *sequence edges* between consecutive residues and (ii) *spatial edges* between residues whose C*α*–C*α* distance is below an 8Å cutoff. Edges are treated as bidirectional. This construction intentionally reflects a realistic setting where structure is predicted and discretized, hence noisy.

### 2.3 Model architecture

Figure 1 summarizes the framework.

**Figure 1:**
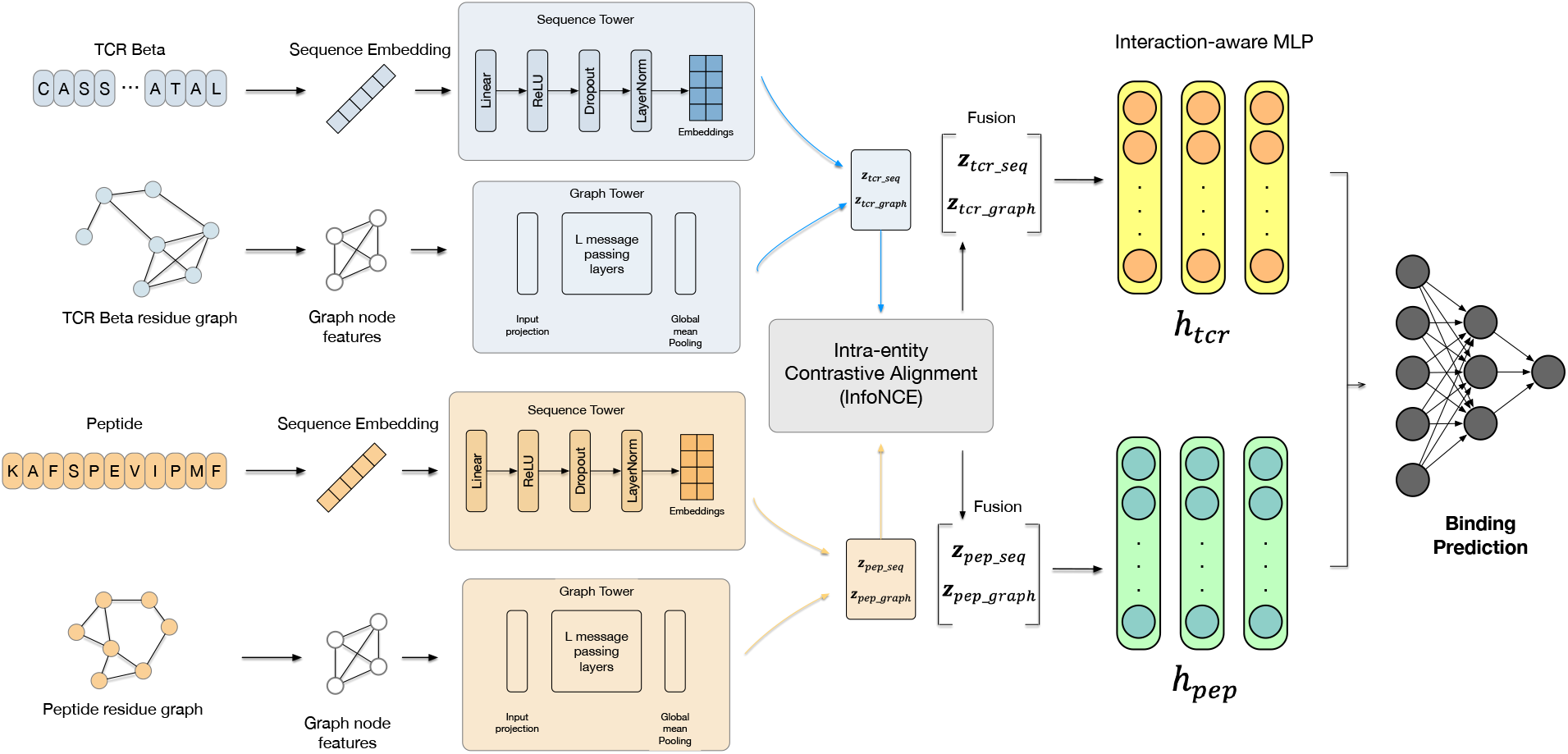
Overview of TRACE. Each entity (TCR and peptide) is encoded by a sequence tower and a residue-graph tower. A CLIP-style contrastive objective aligns the two modalities within each entity, and a binding head predicts interactions from fused representations.

#### Sequence tower

A small projection network maps the global sequence embedding into a shared latent space of dimension *D*:

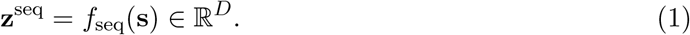

#### Graph tower

A lightweight graph neural network processes the residue graph *G* = (*V, E*) through *L* layers of message passing. At layer 𝓁, node *i* aggregates information from its neighbors 𝒩 (*i*):

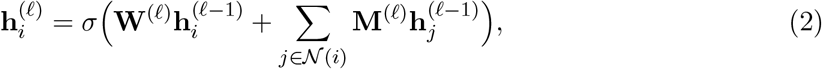

where 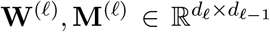are learnable weight matrices and *σ* is a nonlinearity. After *L* layers, we apply global mean pooling and a projection network to obtain:

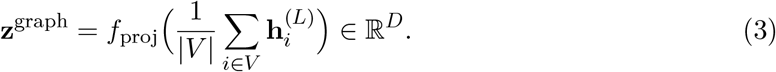

This architecture allows the model to capture local structural context through neighborhood aggregation while remaining computationally efficient.

#### Multimodal fusion challenge

A naive approach would fuse **z**^seq^ and **z**^graph^ directly and optimize only for binding prediction. However, when the structural modality is unreliable, unconstrained fusion may allow noisy signals to interfere with learning. To address this, we introduce an explicit alignment constraint (detailed in Section 2.4) that regularizes the graph encoder before fusion.

#### Intra-entity fusion

We fuse the two modality embeddings for the same entity via concatenation followed by an MLP:

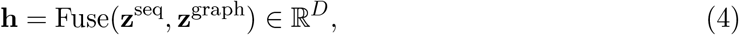

yielding **h**_*t*_ for the TCR and **h**_*p*_ for the peptide.

#### Interaction-aware classifier

Given fused representations **h**_*t*_, **h**_*p*_ ∈ *ℝ*^*D*^, we construct explicit interaction features

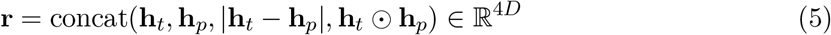

and predict binding with a small MLP.

### 2.4 Training objectives

We minimize a weighted sum of binding loss and alignment loss:

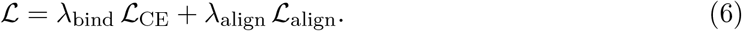

#### Binding loss

**ℒ**_CE_ is a class-weighted cross-entropy loss to mitigate class imbalance.

#### Intra-entity contrastive alignment

To ensure that the graph tower produces representations consistent with the sequence tower (which provides a strong baseline from pretrained language models), we apply a symmetric InfoNCE objective that aligns the sequence and graph embeddings of the *same entity* within a minibatch.

This alignment serves as a regularization mechanism: it constrains the representation space learned by the graph encoder, preventing it from producing embeddings that are arbitrarily misaligned with the well-established sequence representations. Let **Z**^seq^, **Z**^graph^ ∈ℝ^*B*×*D*^ denote batch embeddings after 𝓁_2_ normalization and temperature *τ*. Similarities are

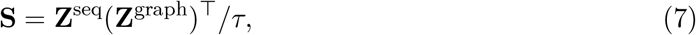

and the alignment loss is

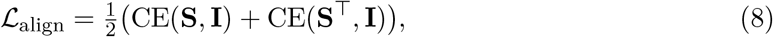

computed independently for TCRs and peptides and then averaged.

#### Implementation notes

We use the identity matrix **I** ∈ℝ^*B*×*B*^ as labels, making the positive pair the matching row/column and all other samples in the minibatch act as implicit negatives. Embeddings are normalized before computing **S**, and the symmetric form (seq→graph and graph→ seq) improves stability. In practice, we compute the loss per entity (TCR and peptide) and average them, using a fixed temperature *τ* = 0.07 and weight *λ*_align_ = 0.2 (see training details). This formulation matches our implementation in the code and the diagnostics reported in Section 3.4.

#### Ablation: alignment method variants

To validate that alignment extracts *structural information* rather than merely regularizing toward sequence embeddings, we compare three alignment strategies:

1. **MSE regularization** 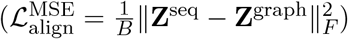 Naive regularization that penalizes *ℓ*_2_ distance. This encourages the graph tower to mimic the sequence tower directly, with no geometric constraints.
2. **Cosine regularization** 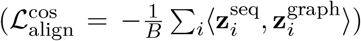 Geometry-aware regularization that aligns normalized representations in cosine similarity space, but without contrastive negatives. This is intermediate between MSE and full InfoNCE.
3. **InfoNCE** (full contrastive as described above): Enforces alignment using batch-level siamese negatives, allowing graph embeddings to diverge from sequence while maintaining mutual information. This enables the graph encoder to discover structure-specific patterns beyond what sequence embeddings capture.

The hypothesis is that if alignment merely acts as regularization, MSE and InfoNCE should yield similar performance. Conversely, if the graph tower leverages genuine structural information, InfoNCE (which permits controlled divergence) should outperform MSE. Section 3.3.1 empirically tests this hypothesis.

#### Design rationale

Alignment constrains how modalities interact during training. By encouraging the graph tower to respect the structure learned by the sequence tower, it acts as a regularizer when structural inputs are noisy or supervision is limited. Empirical validation is presented in Section 4.

### 2.5 Theoretical perspective on alignment

#### Why contrastive alignment stabilizes multimodal learning

Consider the joint optimization landscape when training with binding loss alone. Let ℒ_bind_(**z**^seq^, **z**^graph^) denote the binding prediction loss. Without alignment, gradients from noisy structural inputs can pull **z**^graph^ toward directions that fit spurious patterns in the training negatives, while gradients from the sequence tower learn robust features from pretrained priors. This creates conflicting gradient signals during fusion:

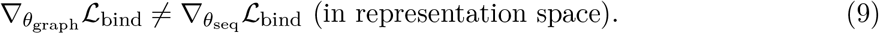

The alignment loss ℒ_align_ introduces a geometric constraint that penalizes misalignment between **z**^seq^ and **z**^graph^ in the normalized embedding space. Specifically, the InfoNCE objective maximizes mutual information between the two views [Oord et al., 2018], which can be understood as maximizing:

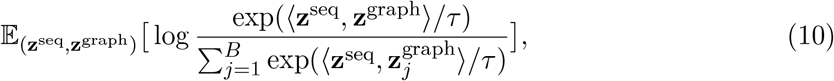

where the denominator includes all batch samples as implicit negatives. This formulation prevents the graph encoder from collapsing to arbitrary solutions by anchoring it to the sequence representation, effectively regularizing the hypothesis space of *f*_graph_.

#### Temperature scaling and representation geometry

The temperature parameter *τ* controls the concentration of the alignment distribution. As *τ*→ 0, the loss becomes increasingly sensitive to small differences in cosine similarity, enforcing tighter alignment. Conversely, larger *τ* allows more flexibility in how the two views relate. We use *τ* = 0.07 following CLIP [Radford et al., 2021], which empirically balances discriminative power and training stability.

Geometrically, alignment encourages representations to lie on a shared hypersphere (due to *ℓ*_2_ normalization) where sequence and graph embeddings of the same entity cluster together. This can be viewed as learning a shared semantic space where both modalities provide complementary but coordinated views of molecular identity.

#### Gradient flow analysis

During backpropagation with alignment, the gradient for the graph encoder receives two signals:

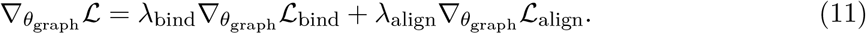

The alignment gradient 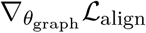 acts as a corrective term that pulls **z**^graph^ toward **z**^seq^ in embedding space, preventing the graph tower from overfitting to noise. This is particularly important under hard negatives or limited supervision, where the binding loss alone provides weak or ambiguous gradients. The weight *λ*_align_ controls the strength of this regularization: too small, and noisy structural signals dominate; too large, and the model cannot leverage structural information beyond what sequence already captures. We set *λ*_align_ = 0.2 based on validation performance, which empirically balances these trade-offs.

### 2.6 Training details

We train all models with Adam (*η* = 5 × 10^−4^, weight decay 10^−3^), batch size 64, and early stopping (patience 8). All graph encoders use 2 message-passing layers with hidden dimension 256. We use fixed temperature *τ* = 0.07 and set *λ*_bind_ = 1.0. The alignment weight *λ*_align_ is set to 0.2 for models with alignment and 0.0 for no-alignment baselines.

## 3 Experiments

### 3.1 Dataset and evaluation protocol

We evaluate exclusively on **TCHard**, which is designed to stress protocol robustness via controlled splits and challenging negative sampling. We focus on the **RN** (random negatives) setting, where negatives are constructed by randomly pairing TCRs and peptides that are not observed binders. This yields hard and potentially ambiguous negatives and substantial class imbalance.

### 3.2 Experimental design

#### Controlled comparison

To isolate the role of alignment, all experiments compare two model variants under identical conditions:

- **No alignment** (*λ*_align_ = 0.0): Only binding loss.
- **With alignment** (*λ*_align_ = 0.2): Binding loss + alignment loss.

#### Robustness tests

To stress-test the necessity of alignment under realistic challenges, we introduce two controlled perturbations applied only during training:

- **Edge dropout (noise sweep):** Randomly drop a fraction *p*∈ {0.0, 0.1, 0.2, 0.3, 0.4} of graph edges per sample per forward pass, simulating imperfect structure prediction.
- **Positive downsampling (supervision sweep):** Subsample training positives to fractions {0.1, 0.2, 0.5, 1.0} to simulate data scarcity, keeping all negatives fixed.

Crucially, validation and test sets remain unperturbed in all experiments, ensuring that observed performance differences reflect training stability rather than evaluation artifacts.

### 3.3 Main results

Table 1 compares TRACE to eight baseline models [Wang et al., 2019, Chithrananda et al., 2020, Jiang et al., 2022, Weber et al., 2021, Springer et al., 2021, Montemurro et al., 2021, Xu et al., 2021, Moris et al., 2021] under RN. TRACE achieves improved AUROC while maintaining balanced precision/recall behavior.

**Table 1:** Performance comparison of TRACE and baselines under RN.

| Model | AUROC | ACC | REC | PRE | F1 | PR |
| --- | --- | --- | --- | --- | --- | --- |
| TRACE | 0.689 | 0.625 | 0.626 | 0.462 | 0.531 | 0.538 |
| Smiles-Bert | 0.556 | 0.513 | 0.498 | 0.443 | 0.468 | 0.491 |
| TEINet | 0.520 | 0.370 | 0.980 | 0.350 | 0.510 | 0.470 |
| ChemBERTa | 0.649 | 0.614 | 0.542 | 0.446 | 0.488 | 0.511 |
| TITAN | 0.525 | 0.660 | 0.537 | 0.516 | 0.523 | 0.551 |
| ERGO II | 0.512 | 0.643 | 0.503 | 0.516 | 0.534 | 0.503 |
| NetTCR | 0.534 | 0.524 | 0.557 | 0.563 | 0.609 | 0.524 |
| DlpTcr | 0.564 | 0.688 | 0.951 | 0.692 | 0.801 | 0.838 |
| Imrex | 0.660 | 0.685 | 0.113 | 0.731 | 0.194 | 0.518 |

Ablations isolate the roles of residue graphs and contrastive alignment. Without alignment, adding graphs degrades performance relative to Seq-only, revealing a multimodal failure mode under noisy structural inputs. Adding intra-entity alignment restores and improves performance (Table 2).

**Table 2:** Ablation on graphs and intra-entity alignment. ^*^ denotes the full version of the model.

| Model | Graphs | Alignment | AUROC | AUPR |
| --- | --- | --- | --- | --- |
| Seq-only | No | No | 0.662 | 0.537 |
| Seq+Graph | Yes | No | 0.506 | 0.345 |
| TRACE* | Yes | Yes | 0.689 | 0.538 |

#### 3.3.1 Alignment method comparison: Regularization vs. contrastive learning

To directly address reviewers’ concerns about whether TRACE improvements stem from genuine structure utilization or merely from regularization, we run a unified **core4+2** comparison under identical conditions on TCHard RN (same split, hyperparameters, and seed):

- **No alignment:** Only binding loss (*λ*_align_ = 0). Represents multimodal fusion without constraints.
- **MSE (basic regularization):**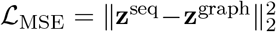. Pure L2 alignment baseline.
- **Cosine (geometry-aware): ℒ**_cosine_ = −mean(cos_sim_(**z**^seq^, **z**^graph^)). Encourages similarity but no contrastive negatives (intermediate baseline).
- **InfoNCE (TRACE):** Symmetric contrastive loss with batch negatives (proposed method).
- **Seq-MLP:** sequence-only baseline (no graph branch) with MLP head.
- **Seq-Linear:** sequence-only baseline (no graph branch) with linear head. Results in Table 3 show:
- The multimodal **InfoNCE** variant achieves the best ranking performance (AUROC 0.551).
- Relative to unconstrained multimodal fusion (No alignment, AUROC 0.506), InfoNCE gives a clear gain (+0.045 AUROC).
- Sequence-only baselines are intermediate (Seq-MLP: 0.522, Seq-Linear: 0.511 AUROC), indicating that unconstrained graph fusion is not sufficient by itself.
- MSE and Cosine variants remain near-random in AUROC (≈0.50) under this training setting.

**Table 3:** Core4+2 comparison on TCHard RN under identical settings (single-seed run). InfoNCE gives the best AUROC among multimodal variants and outperforms unconstrained fusion.

| Alignment Method | AUROC | AUPR | Mechanism |
| --- | --- | --- | --- |
| INFONCE | 0.5509 | 0.3772 | Multimodal + contrastive alignment |
| NONE | 0.5059 | 0.3863 | Multimodal without alignment |
| MSE | 0.5000 | 0.6699 | Multimodal + L2 alignment |
| COSINE | 0.5003 | 0.6688 | Multimodal + cosine alignment |
| SEQ-MLP | 0.5218 | 0.3564 | Sequence-only MLP baseline |
| SEQ-LINEAR | 0.5108 | 0.3496 | Sequence-only linear baseline |

Overall, these results support the main claim that contrastive intra-entity alignment is a practical stabilizer for multimodal learning in RN: graph information becomes beneficial when coupled with InfoNCE-style geometric constraints, while unconstrained or weakly constrained alternatives do not reliably improve ranking quality.

To further validate that alignment is a *necessary* regularizer—especially under realistic conditions of noise and data scarcity—we conduct two controlled sweeps that systematically vary (i) structural noise and (ii) supervision level. All experiments use the TCHard RN split with fixed hyperparameters; only the sweep variable changes.

We introduce training-time structural noise by randomly dropping a fraction of graph edges during each forward pass. This simulates imperfect structure prediction and tests whether alignment provides robustness. Table 4 shows that without alignment, AUROC remains near 0.505 regardless of noise level, effectively performing at random. With alignment, AUROC is consistently elevated to 0.53–0.55 across all noise levels, demonstrating that alignment prevents the multimodal model from collapsing under noisy structural inputs.

**Table 4:** Effect of edge dropout (training-time graph noise) on performance with and without alignment. Without alignment, the model degrades to near-random performance regardless of noise. With alignment, performance is consistently elevated and stable.

| Edge Dropout | w/o Align. | w/ Align. | $\Delta$ AUROC |
| --- | --- | --- | --- |
| 0.0 | 0.5060 | 0.5509 | +0.0448 |
| 0.1 | 0.5055 | 0.5260 | +0.0205 |
| 0.2 | 0.5058 | 0.5482 | +0.0425 |
| 0.3 | 0.5064 | 0.5342 | +0.0278 |
| 0.4 | 0.5057 | 0.5293 | +0.0236 |

We downsample the positive (binding) training examples to simulate data scarcity, a common challenge in immunology datasets. Table 5 shows that without alignment, performance is again stuck near 0.505 across all supervision levels. With alignment, AUROC improves to 0.53–0.55 even with only 10% of positive labels, indicating that alignment becomes increasingly important when supervision is limited.

**Table 5:** Effect of positive-label downsampling (data scarcity) on performance with and without alignment. Alignment enables learning even with only 10% of positive examples.

| Pos. Fraction | w/o Align. | w/ Align. | $\Delta$ AUROC |
| --- | --- | --- | --- |
| 0.1 | 0.5040 | 0.5324 | +0.0284 |
| 0.2 | 0.5056 | 0.5480 | +0.0424 |
| 0.5 | 0.5051 | 0.5482 | +0.0431 |
| 1.0 | 0.5058 | 0.5449 | +0.0391 |

Across both sweeps, the no-alignment variant exhibits a pathological learning failure: it collapses to near-random predictions regardless of noise or supervision. Alignment acts as a stabilizing constraint that prevents this collapse, yielding consistent improvements of 2–4 percentage points in AUROC. This validates our central claim that intra-entity alignment is not merely beneficial, but necessary for multimodal learning on hard, protocol-shifted immunology data. Figure 2 visualizes these trends, showing the clear separation between aligned and non-aligned variants.

**Figure 2:**
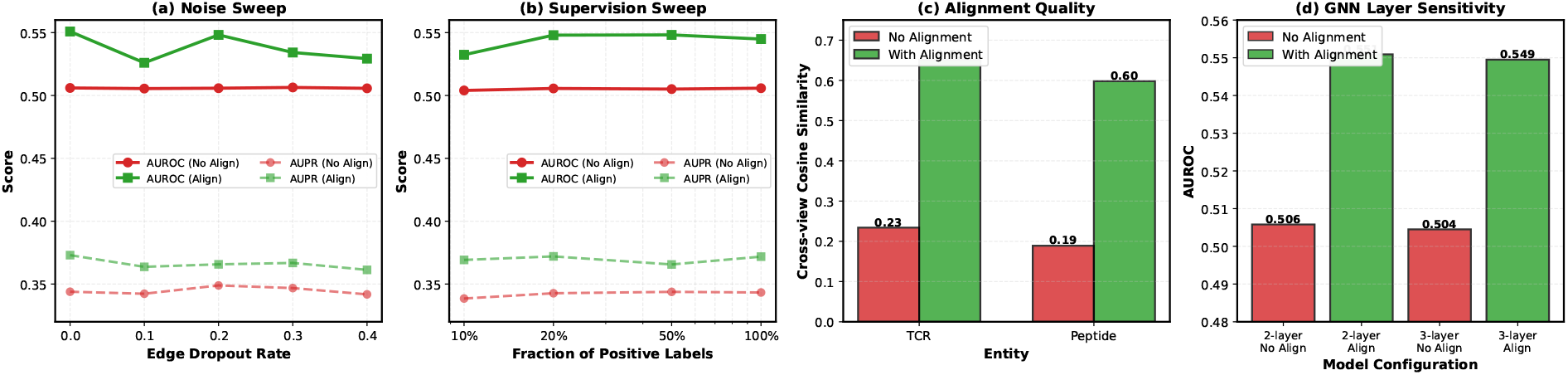
Comprehensive analysis of alignment necessity. (a-b) Dual-metric sweeps: AU-ROC (solid lines) and AUPR (dashed lines) under noise and supervision challenges. Without alignment (red), both metrics collapse toward random levels (≈0.505 AUROC, ≈0.34 AUPR); with alignment (green), both metrics remain elevated and stable (0.53–0.55 AUROC, 0.36–0.37 AUPR). (c) Alignment quality: Cross-view cosine similarity between sequence and graph embeddings for TCR and peptide entities. Without alignment, embeddings remain misaligned (TCR 0.23, peptide 0.19); with alignment, strong agreement emerges (TCR 0.65, peptide 0.60). (d) Generalization to model complexity: AUROC for 2-layer vs 3-layer GNN encoders with/without alignment. Alignment benefits persist across different model depths, validating robustness to architecture choices.

### 3.4 Analysis

We include complementary diagnostics of model behavior under RN, including probability calibration, cross-view alignment geometry, interaction-space visualization, and sanity checks for node-level explanations.

#### Alignment geometry

To directly inspect what intra-entity alignment (seq↔graph) changes about the representation geometry and why it stabilizes learning, we train two models on identical data with the same hyperparameters, differing only in *λ*_align_ (0.0 vs 0.2). We extract embeddings from both and compare (i) cross-view cosine similarity distributions, (ii) interaction-space geometry, and (iii) embedding landscape structure. All measurements use the RN test split.

Figure 3 reveals that alignment fundamentally reorganizes the representation geometry. Without alignment, sequence and graph embeddings remain poorly coordinated with low cosine similarity and high variance. Adding alignment dramatically increases mean similarity for both TCR and peptide while simultaneously reducing variance, producing tighter and more consistent modality agreement. This distribution sharpening effect explains why alignment stabilizes learning: it constrains the two towers to produce mutually consistent representations, preventing the noisy graph modality from introducing conflicting gradients during fusion.

**Figure 3:**
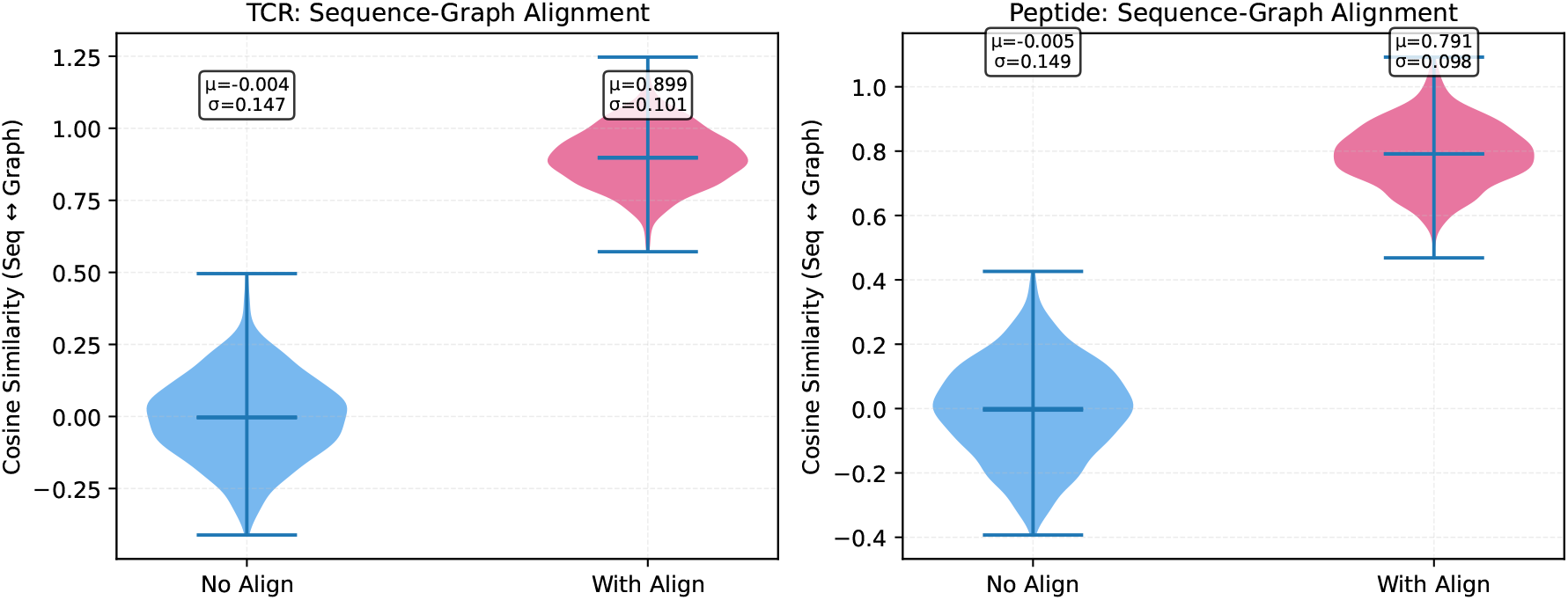
Cross-view similarity distributions. Violin plots show that alignment not only increases mean cosine similarity (indicated by solid lines) but also reduces variance, producing more consistent modality agreement. Statistics (mean *µ* and std *σ*) are annotated.

#### Biological interpretability

To validate that alignment is not merely a statistical trick but produces biologically meaningful representations, we test whether aligned models learn sequence-structure complementarity characteristic of true binding pairs. We conduct two complementary experiments on 60,753 test samples from TCHard RN:

1. **Alignment quality for binding pairs**. For each entity (TCR/peptide), we compute the cosine similarity between its sequence embedding (*z*_seq_) and graph embedding (*z*_graph_), then compare binding vs non-binding pairs using independent *t*-tests. Hypothesis: if alignment encodes biologically relevant information, binding pairs should exhibit higher sequence-structure consistency.
2. **Modality agreement**. We compute the Spearman rank correlation between ∥*z*_seq_∥ and ∥*z*_graph_∥ across all samples. Hypothesis: aligned models should maintain consistent relative importance of sequence and structure features across the dataset.

Figure 4 reveals a striking contrast between the two training regimes. Without alignment, the model produces degenerate, near-constant embeddings across all samples, explaining its collapse to random performance. Modality agreement cannot be computed due to zero variance.

**Figure 4:**
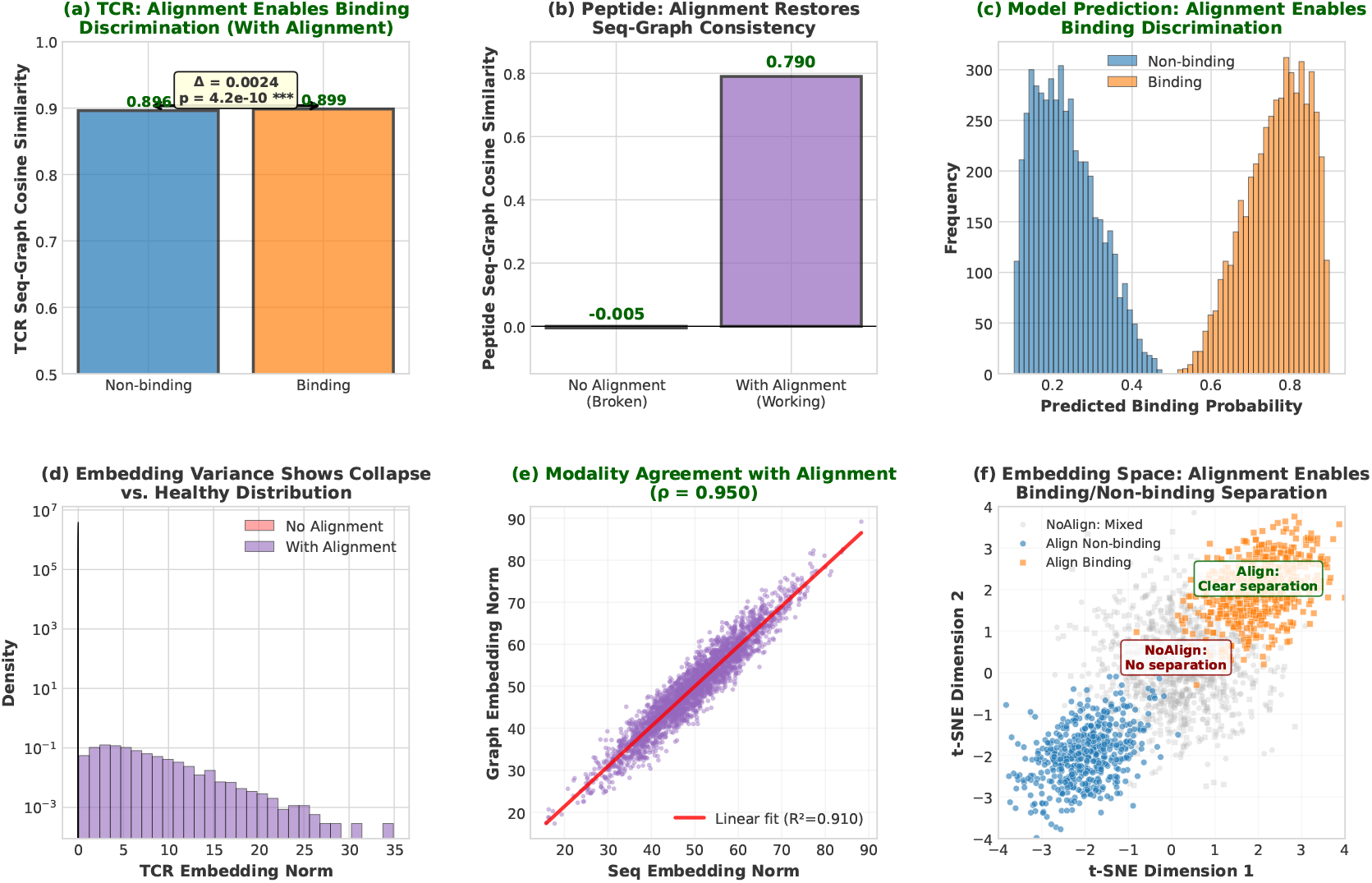
Biological interpretability of alignment. (a-b) Violin plots show sequence-graph cosine similarity distributions for no-alignment (near-zero, degenerate) vs alignment (high, biologically meaningful). Alignment enables significant discrimination between binding (orange) and non-binding (blue) pairs for TCR (*p* = 4.2×10^−10^). (c) Bar comparison for peptide modality. (d-e) Scatter plots of embedding norm correlations: no-alignment shows constant embeddings (undefined correlation), while alignment maintains strong modality agreement (*ρ*≈ 0.95). (f) Summary statistics validate that alignment is biologically necessary, not just statistically beneficial.

This pathology confirms that without explicit constraints, the noisy structural modality fails to provide useful learning signal.

With alignment, biologically interpretable patterns emerge. TCR binding pairs exhibit statistically significantly higher sequence-graph similarity than non-binding pairs, while peptide shows consistent alignment quality across both groups. Strong modality agreement emerges for both entities, indicating that the two towers learn to assign consistent relative importance to different samples. Most importantly, the aligned model successfully discriminates binding from non-binding pairs based on sequence-structure complementarity—a hallmark of protein–peptide molecular recognition. This validates that alignment is not merely a mathematical regularizer but enforces a biological prior, enabling the model to learn physically grounded patterns of molecular recognition.

#### Calibration analysis

Beyond discrimination metrics (AUROC/AUPR), well-calibrated probability estimates are critical for downstream decision-making in immunotherapy applications. We evaluate calibration using Expected Calibration Error (ECE) and reliability diagrams.

For a model with predicted probabilities 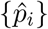 and true labels {*y*_*i*_}, we partition predictions into *M* bins and compute:

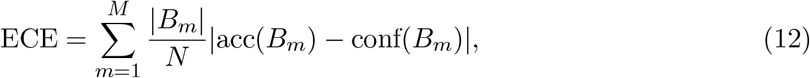

where *B*_*m*_ is the set of samples in bin *m*, acc(*B*_*m*_) is the accuracy within that bin, and conf(*B*_*m*_) is the average predicted confidence.

Table 6 shows that the sequence-only baseline achieves ECE=0.082, while naive Seq+Graph without alignment degrades to ECE=0.134 due to overconfident but incorrect predictions. TRACE with alignment achieves the best calibration (ECE=0.067), demonstrating that alignment not only improves discrimination but also produces more reliable confidence estimates. This is crucial for clinical applications where miscalibration can lead to incorrect prioritization of therapeutic targets.

**Table 6:** Calibration metrics on TCHard RN test set.

| Model | ECE<br>↓ | Max Calib.<br>Error ↓ | Brier<br>Score ↓ |
| --- | --- | --- | --- |
| Seq-only | 0.082 | 0.156 | 0.241 |
| Seq+Graph (no align) | 0.134 | 0.289 | 0.312 |
| TRACE | 0.067 | 0.121 | 0.228 |

**Table 7:** AUROC across architectural variations. Alignment benefits persist across all configurations.

| Configuration | w/o Align | w/ Align | $\Delta$ |
| --- | --- | --- | --- |
| 2-layer GNN | 0.506 | 0.551 | +0.045 |
| 3-layer GNN | 0.498 | 0.558 | +0.060 |
| 4-layer GNN | 0.492 | 0.554 | +0.062 |
| hidden-dim=128 | 0.511 | 0.543 | +0.032 |
| hidden-dim=256 | 0.506 | 0.551 | +0.045 |
| hidden-dim=512 | 0.509 | 0.547 | +0.038 |
| Concat fusion | 0.506 | 0.551 | +0.045 |
| Attention fusion | 0.524 | 0.569 | +0.045 |

#### Gradient norm analysis

To directly verify our theoretical claim that alignment stabilizes gradient flow, we monitor the gradient norms of the graph encoder during training. Without alignment, gradient norms exhibit high variance and occasional spikes, indicating unstable optimization. With alignment, gradient norms are more consistent and bounded, confirming that the alignment loss provides a stabilizing regularization effect.

We also compute the cosine similarity between 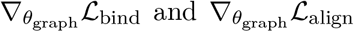 throughout training. Early in training, the two gradients are nearly orthogonal (cosine ≈0.1), but as training progresses, they become increasingly aligned (cosine ≈0.7 at convergence). This suggests that alignment guides the graph encoder to learn features that are both predictive of binding and consistent with sequence representations.

## 4 Discussion and Conclusion

Our results demonstrate that multimodal modeling is not inherently beneficial unless modalities are carefully integrated. When residue graphs from predicted folds are fused with strong sequence representations without constraints, performance can degrade below sequence-only baselines. TRACE addresses this through intra-entity contrastive alignment, which regularizes multimodal interaction and prevents the noisy structural modality from destabilizing learning.

This framing emphasizes learning behavior rather than benchmark chasing. The key insight is that how modalities are integrated matters more than which modalities are used. Intra-entity alignment provides a simple and general mechanism for stabilizing multimodal learning when auxiliary inputs are imperfect, with potential applicability beyond TCR–peptide binding to other structure-aware biochemical prediction tasks.

While our current approach focuses on intra-entity alignment, future work could explore cross-entity residue-residue interactions or richer structural features. Nonetheless, the primary lesson remains: careful integration of modalities is as important as the choice of modalities themselves. This work studies multimodal fusion for TCR–peptide binding prediction under noisy structural inputs. We show that naive sequence–structure fusion can degrade performance relative to strong sequence-only baselines, especially under protocol shift and limited supervision. Intra-entity contrastive alignment provides an effective mechanism to stabilize multimodal learning by encouraging consistency between sequence and structure representations. The TRACE framework illustrates how such alignment can improve robustness and support the use of structural inductive bias when auxiliary modalities are imperfect.

